# Temporal autocorrelation increases temperature-driven extinction risk by clustering stressful conditions

**DOI:** 10.1101/2025.07.29.667527

**Authors:** Alison J. Robey, Misha T. Kummel, David A. Vasseur

## Abstract

Environments are becoming increasingly autocorrelated as global climate change progresses, leading to the intensification of deadly events like heatwaves and droughts. Theory shows that temporally-autocorrelated environments generate a higher risk of population extinction; however, little work has been done to incorporate temporal autocorrelation into thermal performance-based projections of extinction risk. Here, we pair stochastic simulation models of population dynamics with systematically generated temperature time series to determine when higher levels of autocorrelation generate greater extinction risk. We show that autocorrelation is a significant mediator of risk under stressful temperature regimes, with important ramifications for forecasting temperature-driven extinctions in ectothermic organisms. We validate our predictions with a factorial experiment in microcosms of the single-celled protist *Paramecium caudatum*. Taken together, these results provide the foundation for predicting which species and environments face the greatest risks under increasing autocorrelation.

## 1 Introduction

Anthropogenic climate change is reshaping both the magnitude and the structure of environmental temperature variation. While the adverse impacts of increasing thermal means and extremes on species persistence are broadly recognized (e.g., 1), the effects of changing temporal autocorrelation have generally been overlooked. Positive autocorrelation, a positive correlation between proximate values in a sequence, results in the clustering of similar conditions. Increasing levels of autocorrelation thus generate a higher likelihood of remaining near any given temperature for a longer period of time, corresponding to environmental phenomena like longer and more frequent heatwaves. Heatwaves are increasingly responsible for mass mortality events (2), such as the death of 4 million common murres (50% of the population) off the coast of Alaska between 2014 and 2016 (3). Anthropogenic climate change is intensifying heatwave regimes (4) and increasing the autocorrelation of temperatures in many locations across the globe (5; 6), highlighting the urgent need to better understand how the risk of extinction for different species and in different environments changes with the temporal autocorrelation of temperatures.

High temporal autocorrelation affects risk by increasing the propensity for population dynamics to track environmental conditions. In the absence of autocorrelation, successive environmental conditions are unrelated and, if the environment changes frequently enough, population size tends to average across the experienced conditions. When the average environmental conditions are generally favorable, it is thus likely that the population will persist through time (7). In temporally autocorrelated environments, however, successive environmental conditions are correlated and population dynamics track fluctuations more closely (8; 9; 10). If the environment includes a mixture of favorable and unfavorable conditions, the unfavorable conditions are likely to be sequenced together (11), which may cause the population size to approach zero (risking extinction) even when the long-term average is positive (signaling persistence). Autocorrelation thus reduces the predictive power of the mean environmental conditions. Most predictions of temperature-induced extinction risk, however, have focused solely on the (mis)match between organismal thermal tolerance, measured with thermal performance curves (TPCs), and projected changes to thermal distributions (e.g., 12; 13; 14); such forecasting ignores how increasing autocorrelation can increase extinction risk even when thermal means and variances remain unchanged (15).

However, when and how to include changing levels of autocorrelation in risk forecasting remains an open question. Whether environmental autocorrelation should increase extinction risk has been debated because different underlying assumptions produce contradictory outcomes: models where extinction is driven by series of unfavorable conditions (e.g., 16; 17) find that autocorrelation increases risk by increasing the similarity of successive values, while those where extinction can be driven by a single extremely catastrophic event or highly over-compensatory density dependence (e.g., 18) find that autocorrelation decreases risk by reducing the chances of experiencing any catastrophes (19). No previous models, however, deal directly with thermal performance; environmental noise has generally been incorporated into population dynamics as a direct proxy for carrying capacity (e.g., 20) or an additive effect on the population growth rate (e.g., 21). Embedding temperature variation into models of population dynamics is more complex because temperature maps onto one or more parameters via non-linear, unimodal relationships (TPCs; 22). Incorporating these relationships in a biologically meaningful way is a challenge that only very recent work (e.g., 6; 15; 23; 24; 25) has begun to address, and more experimental work is needed to ground-truth the model assumptions underlying such theoretical predictions.

In this project, we pair predictions from stochastic population models with a factorial experiment in microcosms of the single-celled protist *Paramecium caudatum* to assess whether populations experiencing environments with the same thermal distribution but different levels of temporal autocorrelation actually demonstrate differential extinction risk. Previous experimental work with protist microcosms have successfully demonstrated how noise colouration alters population dynamics (26; 27; 28), indicating that such systems are well-suited for decoding the impacts of autocorrelation on persistence. Across theoretical and empirical results, we demonstrate that the mean and temporal autocorrelation of temperature affect extinction risk in tandem. These results provide key insights about how and when risk forecasting efforts need to consider changes to autocorrelation regimes in order to accurately predict extinction events.

## 2 Materials and methods

### 2.1 Population dynamic modeling

Predicting how temporal autocorrelation affects extinction risk requires pairing a model that incorporates temperature into population dynamics with a method to generate autocorrelated temperature time series. Following recent work (6; 9; 15; 24), we use a temperature-dependent version of the *r*-*α* logistic growth model (29). The population growth rate (or Malthusian fitness) *r* is given by a temperature-dependent function *r*(*T*) (a thermal performance curve), while the *α* parameter determines the strength of intraspecific density dependence:

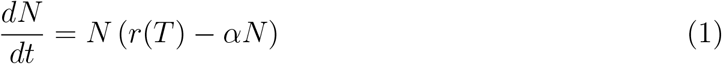

When the thermal environment is favorable (*r*(*T*) *>* 0), the globally stable equilibrium population size (or carrying capacity *K*) is given by *r*(*T*)*/α*. In contrast, when the thermal environment is unfavorable (*r*(*T*) *<* 0), the globally stable equilibrium population size is zero, making the *r*(*T*)-*α* model well-suited for studying extinction risk in thermally variable environments. In this formulation, *α* is assumed to be temperature independent, such that the temperature dependence of *r* and *K* are perfectly correlated (9); while some variation on this pattern is likely (e.g., 30; 31) (and other patterns are possible; see (32; 33)), we retain this assumption for model tractability.

To incorporate temperature variation, we discretize the continuous time model given by Eq. 1 using Euler’s method:

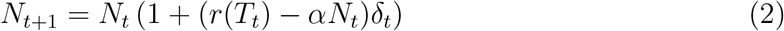

where *r*(*T*_*t*_)−*αN*_*t*_ gives the per-capita rate of change of the population size over the time step *δ*_*t*_ (which is chosen to be very short relative to 1*/r*, the population’s mean generation time). For every time step, the environmental temperature *T*_*t*_ is randomly selected from the normal distribution 𝒩(*µ*_*T*_, *σ*_*T*_), but organized into a time series exhibiting ‘1*/f*^*γ*^-noise’ (20; 34; 35). In 1*/f*^*γ*^-noise, the temporal autocorrelation level is given by the spectral exponent *γ*, where *γ* is the slope of the relationship between the squared amplitude of variability and its frequency *f* in a time series on a log-log scale (e.g., Fig. 2c). Most environmental variables are well-characterized as having positive autocorrelation (*γ >* 0) (36; 37; 38; 39; 40; 41), which is often referred to as ‘red’ noise due to the similar relative importance of low frequencies in red light. Red noise encompasses the range 0 *< γ* ≤ 2, with autocorrelation increasing from white (*γ* = 0) to pink (*γ* = 1) to brown noise (*γ* = 2, consistent with Brownian motion) as the spectral exponent increases (see (42) for a more detailed explanation).

To create temperature sequences with a 1*/f*^*γ*^ structure, we use the method of spectral synthesis (20; 43). Spectral synthesis generates time series with precisely controlled autocorrelation levels for any given mean, variance, and length (though higher moments may vary across replicates when the time series is short and *γ* is large). Other processes for generating autocorrelated time series – such as the continuous-time Ornstein-Uhlenbeck model used previously (24) or its discrete-time equivalent, a first-order autoregressive (AR(1)) model (44) – can only generate noise through a reversion parameter and thus provide much less precise control over the resulting power spectra (40; 43). Additionally, these methods only exhibit 1*/f*^*γ*^-scaling over a certain range of timescales, often with a characteristic loss of power at low frequencies that may not be indicative of natural temperature variation (20; 40; 42).

For any given *γ*, we generate a time series by summing *f* = 1, …, *t*_max_*/*2 sine waves at every *T*_*t*_, where *t* = 1, 2, …, *t*_max_:

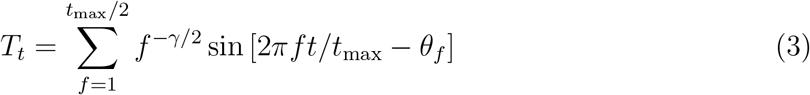

The amplitude *f* ^*−γ/*2^ generates the desired power-law scaling and *θ*_*f*_, a random vector of phase operators drawn from the uniform distribution [0, 2*π*), introduces randomness with-out altering the level of autocorrelation (43). Efficient calculation of the above sum is best computed using an inverse fast Fourier-Transform (45, implemented herein using *Mathemat-ica*’s ‘InverseFourier’ function).

Using the autocorrelated *r*(*T*)-*α* model (Eq. 2, 3), we simulate population dynamics through temperature time series with means ranging from *µ*_*T*_ = [10, 35], standard deviations ranging from *σ*_*T*_ = [0, 8], and levels of temporal autocorrelation ranging from *γ* = [0, 2], assuming a density-dependence parameter *α* = 0.0001 (corresponding with experimental observations that carrying capacity *K* ≈ 5000 individuals) and a time step length *δ*_*t*_ = 0.01. We run 100 replicates per simulation for *t*_max_ =10,000 time steps each, assuming an extinction has occurred if the population size drops below 1 individual. We then compare the simulated extinction risk across autocorrelation levels with two different metrics.

First, we compare the results to the ‘persistence boundaries’ derived in Vasseur et al. (24). Based on classical theory (46; 47), they show that extinction risk can be predicted

for the temperature-dependent *r*(*T*)-*α* model by relying upon analytical derivations of a simpler model, where variation in *r* is normally distributed (with mean 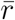 and variance 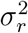) and uncorrelated in time. This work demonstrates that critical changes in the probability density of population size occur when 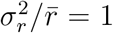 (weak persistence boundary) and 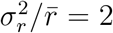 (strong persistence boundary). These boundaries can be traced through the environmental parameter space (gray and black lines, respectively, in Fig. 1a), such that when this ratio is below one (pale yellow region), extended periods of stress – and therefore extinction events – are unlikely; in the intermediate, extinctions become more likely (turquoise region); and above two, extinctions are nearly certain (dark blue region). Although perfect risk predictions are not expected due to the introduction of skewness and higher moments when variation in *T* is filtered into *r* through a non-linear TPC (violating model assumptions), we expect that these boundaries will closely align with simulated results under white noise scenarios but will overpredict persistence under higher autocorrelation.

**Figure 1:**
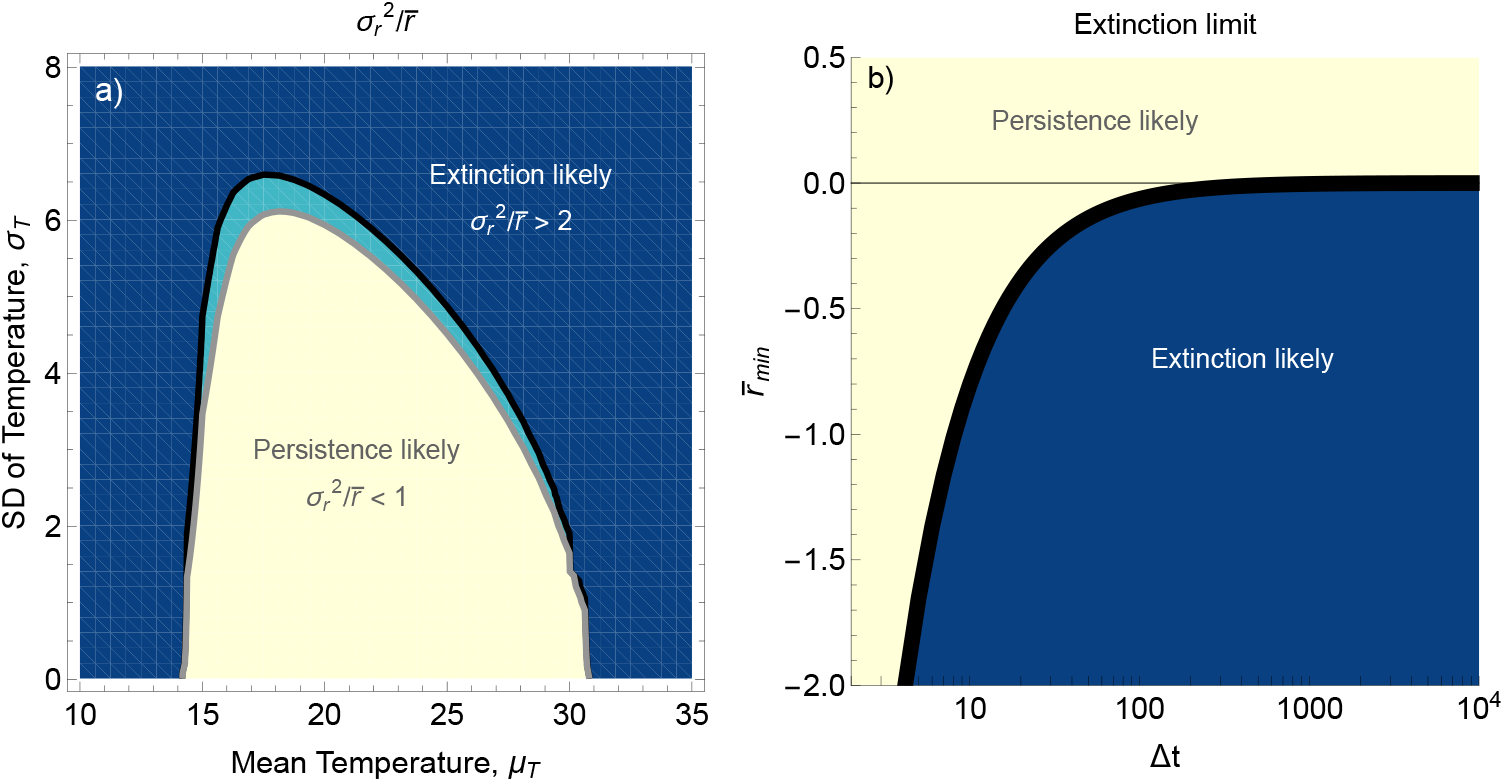
Expected extinction outcomes using (a) the persistence boundaries derived from the ratio of the variance to the mean of fitness 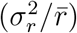 across the temperature parameter space or (b) the minimum mean fitness 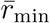 observed across a time window of length Δ*t*, derived from solving the *r*-*α* model for 1 individual given a negative growth rate and a population at carrying capacity (*K* = *N*_0_ = 5000). Shading indicates expected long-term persistence (pale yellow), short-term persistence (turquoise), or extinction (dark blue).

Second, we calculate an extinction boundary based on the temperature sequences themselves. For any stressful temperature event of a given duration (defined as a time period Δ*t* containing *t* time steps where the mean fitness is negative), we calculate a critical value of *r* that delineates how many time steps a population starting from a large size would survive at a constant temperature inducing that mean fitness (i.e., given that the population is shrinking from size *K* at a specific *r <* 0, will it survive for more time steps than Δ*t*?). Above this extinction boundary (black line in Fig. 1b), we expect persistence (pale yellow), and below it, extinction (dark blue). For individual time series with a given *γ*, we then take the minimum average fitness 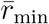 found across the entire series for each interval of length Δ*t*; if those values fall below the critical extinction boundary, extinction is expected. As 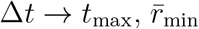 will converge to the average fitness of the entire time series regardless of the ordering (notably, a classic metric of whether populations will persist is whether the value that 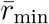 converges to – the average fitness of the entire time series – is positive). However, we expect 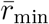 to converge faster for lower levels of autocorrelation, because lower autocorrelation indicates that extremely stressful temperatures are less likely to occur close together. By the same logic, we expect 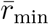 to remain lower under high levels of autocorrelation (even for comparatively longer windows of Δ*t*) because increasing autocorrelation will cause the most stressful temperatures in the time series to clump together. When 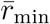 is less than the critical value of *r* that delineates the extinction boundary, we expect that the population has undergone at least one temperature event too stressful to enable persistence.

### 2.2 Thermal Performance Curves

We studied extinction risk in the single-celled protist *P. caudatum* using the same TPC measured and reported in Vasseur et al. (24), collected following the protocol in Wieczynski et al. (23). We counted initial and final population sizes (*N*_0_ and *N*_final_) of 12 replicates grown at 6 constant temperatures (18, 22, 24, 26, 28 and 30^*o*^C) in Percival I-30VL incubators to calculate the population growth rate *r* at each temperature. Each replicate was initialized in a 24-well plate with 4 to 8 protist cells in 1.5 mL of bacterized medium, then incubated for 42 to 68 hours and recounted. We calculated the temperature-dependent population growth rate *r*(*T*) (Fig. 2a) with Eq. 4 and used the ‘*rTPC* ‘package (48) in *R* v4.4.3 (*R* Core Team, 2025) to fit data to the ‘lactin2’ TPC model (49) (Eq. 5):

**Figure 2:**
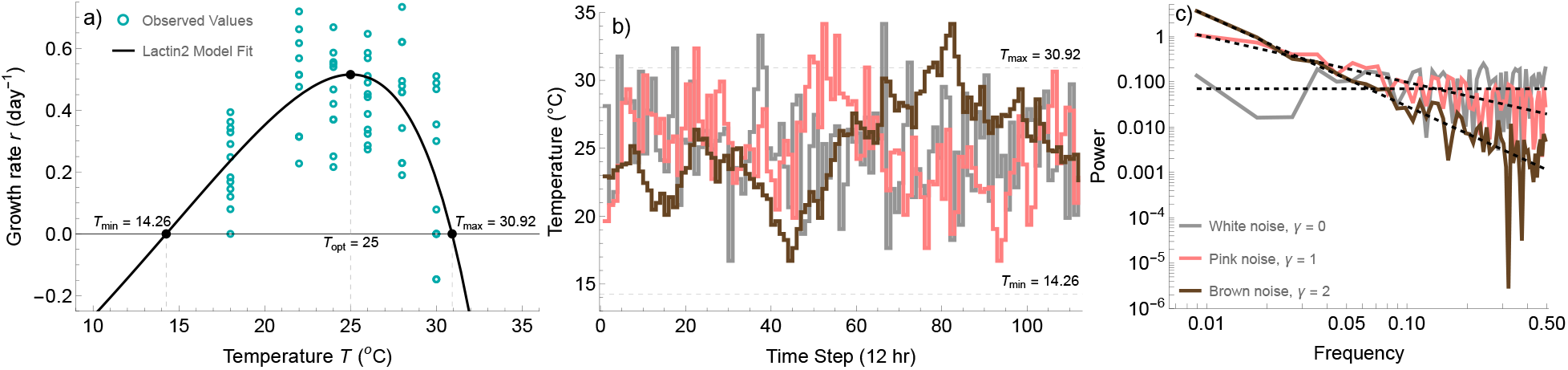
The (a) ‘lactin2’ TPC model fit to experimental data (12 points per temperature) from *P. caudatum*, alongside (b) the three experimental temperature time series, each a permutation of 112 temperatures selected from a normal distribution centered on the TPC’s *T*_opt_ and (c) their spectral densities.

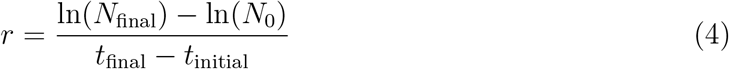

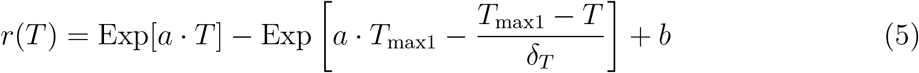

While other TPC models fit the data similarly, this model was chosen because it produces negative growth rates under stressfully hot or cold temperatures (whereas many other models are bounded below by 0, which is inappropriate for a TPC of population growth rate) and is continuously differentiable, which is necessary to calculate persistence boundaries using the numerical expectations of the function’s moments given a normal temperature distribution.

Fitted parameter values were *a* = 0.044, *b* = −1.774, *T*_max1_ = 35.254, and *δ*_*T*_ = 5.435 (see supplementary material S4); approximate TPC parameters are estimated at *T*_min_ = 14.26, *T*_max_ = 30.92, and *T*_opt_ = 25 (where *T*_min_ and *T*_max_ represent the intercepts and *T*_opt_ the peak).

### 2.3 Experimental time series

For experimental treatments, we created three temperature time series with identical distributions but different levels of autocorrelation. First, we created a ‘transfer sequence’ by taking an inverse sample of *t*_max_ = 112 temperatures from the cumulative distribution function of 𝒩(25, 3.5), uniformly spaced such that the discrete temperature distribution used near-perfectly matched the intended normal distribution. We then created ‘reference sequences’ on the interior interval (0, 1) with length *t*_max_ and autocorrelation levels *γ* = 0, 1, 2 using spectral synthesis. Finally, we permuted the values sampled from the distribution 𝒩(25, 3.5) to match the ordinal ranking of the reference sequence. This method, known as spectral mimicry (26), learns the rank ordering of the reference sequence created by spectral synthesis and applies it to the transfer sequence, such that resulting treatments have varying levels of autocorrelation but are composed of the exact same 112 temperature values. We selected three time series corresponding with white (*γ* = 0), pink (*γ* = 1), and brown (*γ* = 2) noise (Fig. 2b); series were generated and spectral exponents calculated (Fig. 2c) using an inverse Fourier transform. Finally, we ran 40 replicates of the *r*(*T*)-*α* model through each selected time series using a stochastic simulation algorithm (SSA; see supplemental material S1) with each time step lasting 12 hours, assuming an initial population size of *N*_0_ = 500, to compare with experimental results.

### 2.4 Extinction experiments

We began experiments by inoculating 210 14 mL culture tubes with 1 mL of stock culture medium containing the bacterivorous protist *P. caudatum* (Fig. 3a), 10 mL of bacterized medium, and one autoclaved wheat seed (Fig. 3b). Bacterized medium was created by autoclaving 1 g of Timothy Hay per 1 L of spring water (obtained from a natural spring in Roxbury, CT, 41°31’07.7”N, 73°15’45.2”W) in a cloth tea bag, then removing hay after 24 hours and adding three bacteria species (*Bactillus subtilis, Escherichia coli* K12, and *Serratia marcescens*), which grew for four days before inoculation. The *P. caudatum* stock culture (containing mentioned bacteria and *Enterobacter aerogenes*) was stored in a constant temperature Percival I-36LL incubator at 22^*o*^C; at inoculation, cell density was approximately 5-15 cells per 0.25 mL. Tubes were then stored with dual position aspirating caps placed in a loose position (to allow gas exchange but avoid contamination) at *T*_opt_ = 25^*o*^C in a constant temperature incubator to stimulate population growth. All consumables, bacteria, and protists were obtained from Carolina Biological Supply.

**Figure 3:**
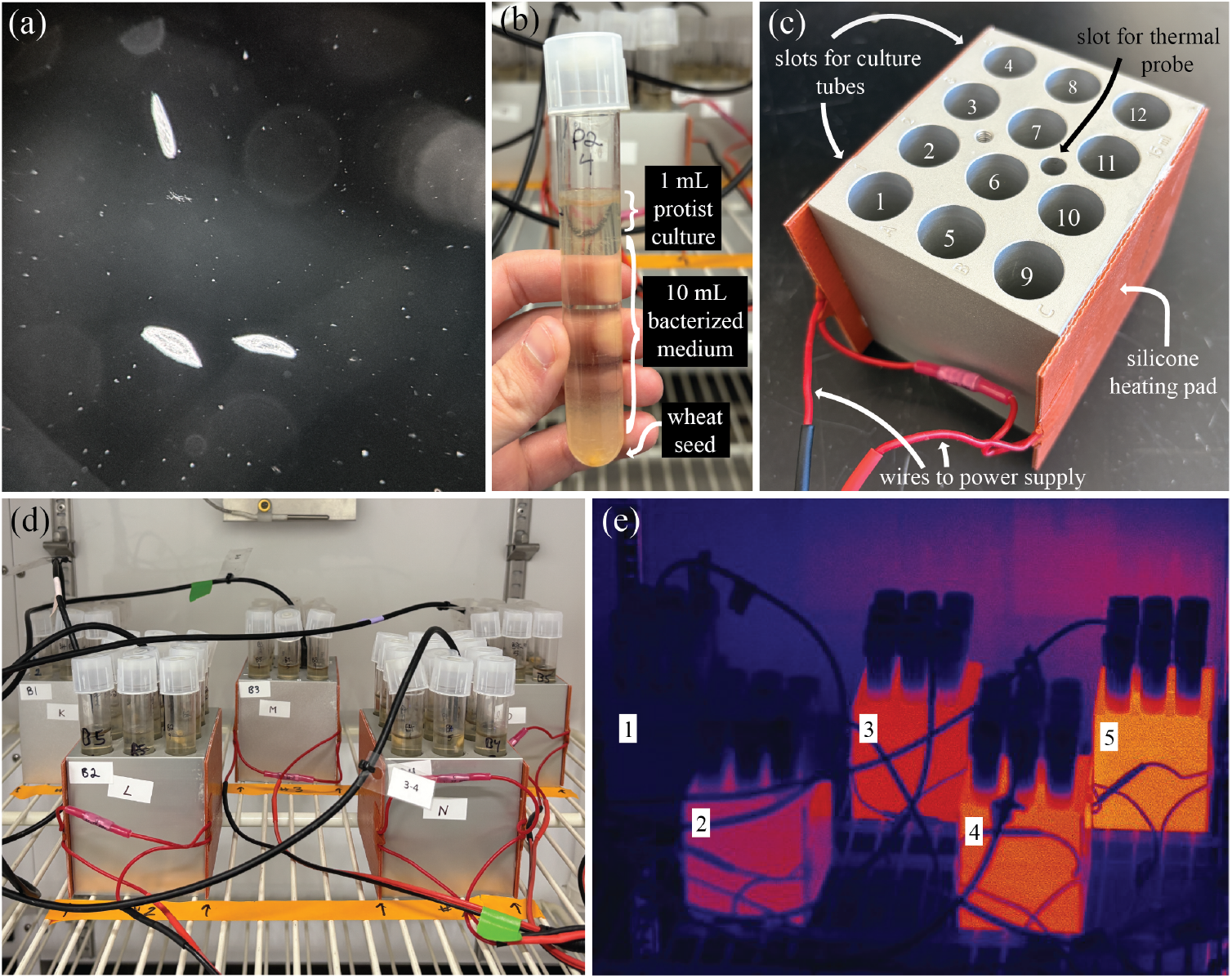
Experimental set-up. (a) Microscope image of 3 *Paramecium caudatum* cells. (b) Example replicate tube with cap in loose position. (c) Example “chilling” block with 12 culture tube slots, a thermal probe slot, a silicone heating pad adhered to either side, and wires connecting them to the power supply. (d) Example incubator set-up with 5 blocks (each at a different mean temperature) per autocorrelation treatment. (e) Thermal image of incubator set-up with heaters activated, showing the thermal gradient across the blocks (brighter colours indicate warmer temperatures).

After one week, we measured cell densities by inverting each tube twice, then extracting 0.25 mL samples with a sterile pipette and manually counting the number of living protists with Leica M125C stereo dissecting microscopes. Prior to the experiment start, 5 failed inoculations were observed and discarded, leaving 205 tubes with persisting populations. The 25 tubes with the lowest population sizes were discarded so that the remaining 180 tubes all contained verified viable populations before the experiment begin (average of 68 cells/mL). Tubes were randomly assigned to treatments such that no treatment began with a significantly different average density.

Tubes were placed into 15 aluminum “chilling” blocks (Research Products International) holding 12 tubes each (Fig. 3c). 5 blocks each were placed into one of three programmable Percival I-30VL incubators (Fig. 3d). Each incubator was outfitted with Intellus Ultra Controllers (Revision 23.01.130, 2.01.00) and programmed with one of the three temperature time series (white, pink, or brown noise; Fig. 2b). Each time step lasted 12 hours; the resulting time series of 112 temperatures thus lasted 56 days. We created a gradient of mean temperature treatments within each incubator by connecting 4 of the 5 blocks to silicone electric heating pads wired to variable DC power supplies; heaters were calibrated to achieve a goal temperature 2^*o*^, 3^*o*^, 4^*o*^, or 5^*o*^ above an ambient of 25^*o*^C before experiment start. The unheated block thus experienced the ambient temperature of the incubator, while each subsequent block connected to a higher voltage was warmed to a set temperature above that ambient (Fig. 3e). Each block was additionally connected to a thermal probe to constantly monitor and record its real-time temperature. We obtained extremely close coupling between the goal and observed block temperatures throughout the experimental time series and across all treatments (Fig. S3, Tables S1 and S2).

Tubes were labeled according their autocorrelation treatment (W = white noise, P = pink noise, B = brown noise), mean temperature treatment (1 = 25^*o*^, 2 = 27^*o*^, 3 = 28^*o*^, 4 = 29^*o*^, 5 = 30^*o*^), and replicate number (1-12), such that, for example, the back left tube experiencing a mean temperature 2^*o*^ above ambient in the pink noise treatment was labeled P2-4 (Fig. 3b). Once a week, every tube was briefly removed from the incubator, topped back up to 11 mL with sterile Timothy Hay medium to compensate for evaporation, and inverted twice to mix in the new medium, facilitate gas exchange, and break up bacterial mats. After 28 and 56 days, we counted cell culture densities using the same methodology described above. If zero cells were observed in the first 0.25 mL sample, we took a second sample; if zero cells were observed in the second sample (indicating a cell density of *<* 2 individuals/mL), we assumed the culture was extinct. After 56 days, we re-collected TPC data from 6 protist populations (using the same methodology as above, but with fewer replicates) for a rough assessment of any significant changes in thermal tolerance. All samples were then disposed.

### 2.5 Statistical analysis

We tested the effects of the mean and autocorrelation of temperature on extinction risk using a bias-reduced binomial generalized linear model (‘brglm’ package (50) in *R* v4.4.3) with mean temperature as a continuous predictor, autocorrelation as a categorical predictor, and extinction as a binary response variable; we also tested for statistically significant effects of initial population density and tube number. This package uses bias-reduction methods to reduce the small-sample bias in maximum likelihood estimates arising from data separation (51), necessary here due to extreme outcomes like perfect prediction of complete extinction or persistence. We assessed model fit based on the Akaike Information Criterion (AIC) scores and comparing residual with null deviance. We additionally tested the significance of binary predictors (parameters located inside versus outside persistence boundaries; tube location on interior versus edges of blocks) using *χ*^2^ tests.

## 3 Results

### 3.1 Modeling results

Using the autocorrelated *r*(*T*)-*α* model, we found that increasing autocorrelation shrinks the envelope of likely persistence by increasing the likelihood that stressful events occur close together. Simulation results are closely predicted by the persistence boundaries derived from the ratio 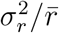 under white noise (Fig. 4a) and under low levels of autocorrelation (*γ <* 0.75; Fig. S1). However, higher levels of autocorrelation result in populations that cannot tolerate as much variability: the parameter space allowing for persistence shrinks rapidly between 0.75 *< γ <* 1, declines marginally and becomes more symmetrical as *γ* → 1.5, then increases very slightly as *γ* → 2 (Fig. 4b, c, S1). Fig. 4d shows the calculated 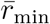 across 100 time series per autocorrelation level for the thermal distribution (25, 3.5) (corresponding to the coloured boxes in panels a-c). Simulations for these parameters found 0%, 85%, and 78% extinction risk under white, pink, and brown noise, respectively, which closely compared to the 3%, 98%, and 79% risk found using the 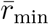 metric. The average 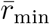 for white noise never dips below the extinction boundary (cyan line compared to black line in Fig. 4d), indicating that the most stressful temperature sequences within those time series are not expected to cause extinction events (particularly if they occur when the population size is large). For pink and brown noise, on the other hand, the average 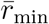 is usually below that extinction boundary, indicating that the most stressful sequences of temperatures in those time series could generate extinction events, even if the population was at carrying capacity when they occurred. Interestingly, while the average 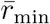 for pink noise is universally less than for white noise, the average for brown noise is greater than either for short Δ*t*s (*<* 7 for white and *<* 100 for pink) due to significantly wider variability between the 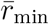 values of different replicates as autocorrelation increases.

**Figure 4:**
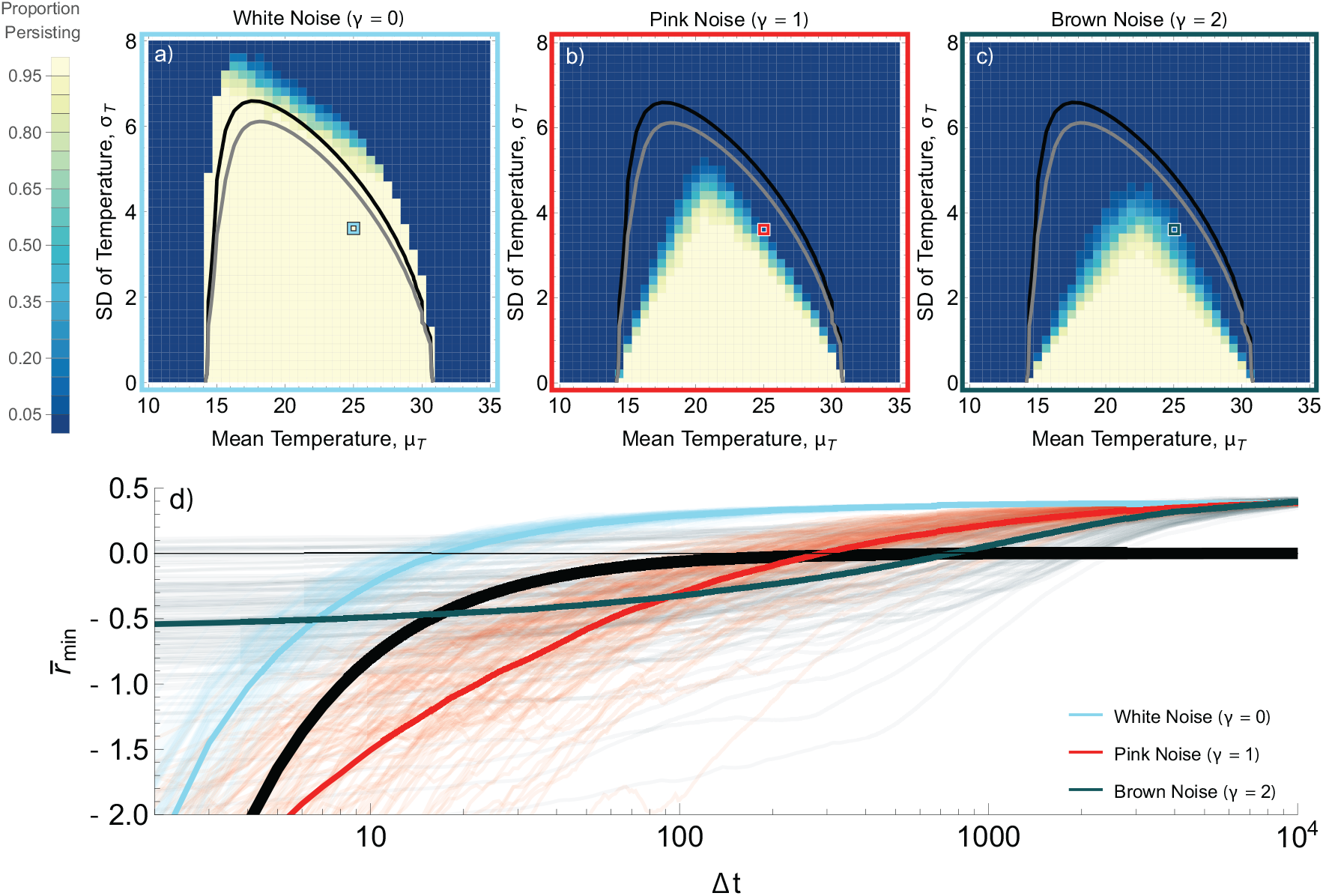
Comparison of the strong (black) and weak (gray) persistence boundaries with the proportion of persisting *P. caudatum* populations under (a) no autocorrelation (white noise, *γ* = 0), (b) medium autocorrelation (pink noise, *γ* = 1), and (c) high autocorrelation (brown noise, *γ* = 2). Each point shows the average of 100 simulations of length *t*_max_ = *α*^*−*1^ = 10000, corresponding with the numerical approximation of expected time to extinction at the weak boundary (24). (d) The minimum growth rates 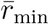 observed for discrete time intervals of length Δ*t* under each level of autocorrelation, with cyan, red, and teal lines showing the average of the 100 faded runs in the same colours, compared to the number of time steps a population can persist with that growth rate from its equilibrium size (thick black line) for the thermal distribution 𝒩 (25, 3.5) (corresponding to the outlined squares in a-c).

### 3.2 Experimental results

Universally, extinction risk increased and population density decreased with mean temperature, with most replicates persisting at the coolest mean temperature (25^*o*^) and all replicates extinct at the hottest mean temperature (30^*o*^; Fig. 5). Consistent with the timing of heat events (where *T > T*_max_), most extinctions in the brown noise treatment occurred between 28 and 56 days, whereas extinctions in the white and pink noise treatments occurred more frequently prior to 28 days (Fig. S4, S5). At 28 days, average population density was highest under brown noise across mean temperature treatments, followed by white noise, then pink noise; at 56 days, average population density was highest for brown, then pink, then white noise at the coolest mean temperature, but lowest for brown noise at all other mean temperatures (Fig. S6). TPCs gathered post-experiment did not show large differences between thermal tolerances of different surviving treatments from either each other or the stock culture, with the exception that the coolest brown noise treatment exhibited noticeably higher growth rates across tested temperatures (Fig. S7).

**Figure 5:**
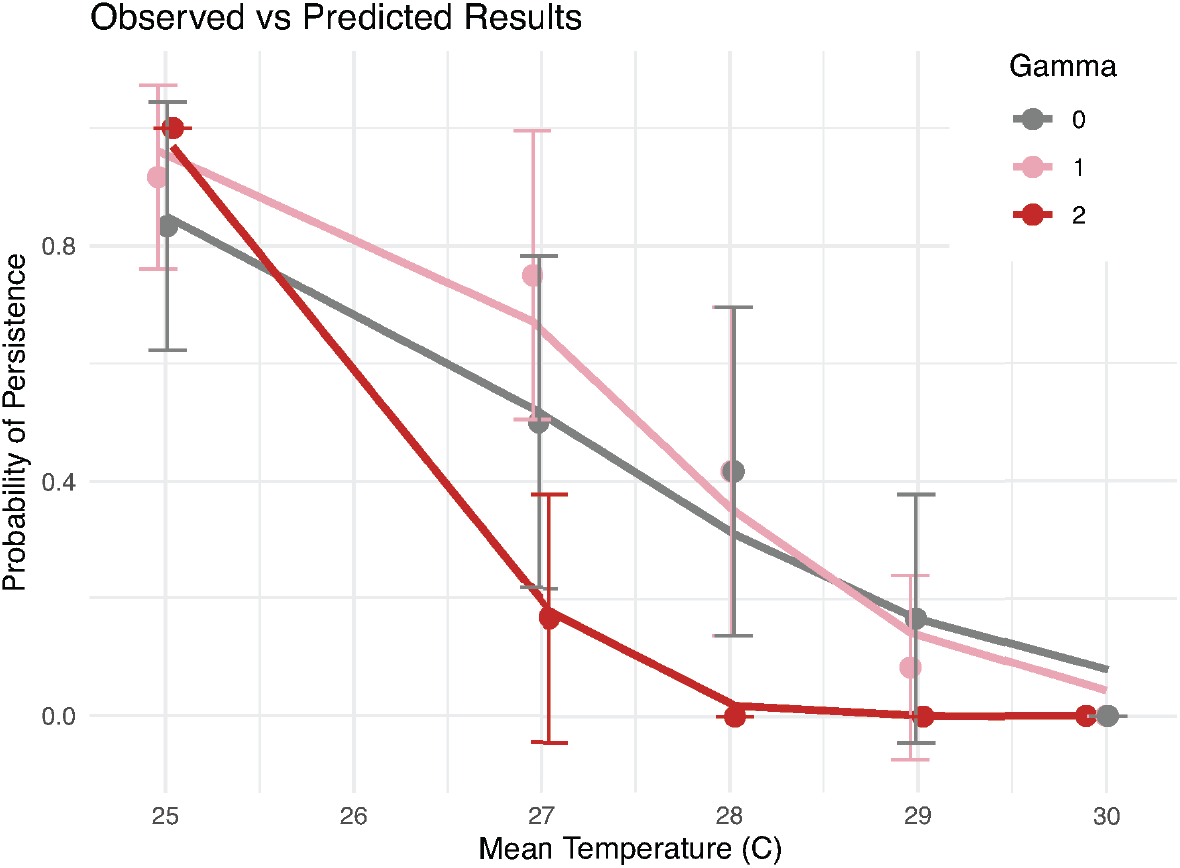
The observed (dots, plotted with 95% confidence intervals) compared to the best model fit prediction (Persistence ∼*µ*_*T*_ ∗*γ*, lines) of persistence probability at each autocorrelation level. Observed points are plotted at the exact mean temperature recorded over the course of the experiment; points obscured by overlap (white and pink noise at 28^*o*^ and 30^*o*^) are identical.

The best fit model for our results was a bias-reduced logistic regression including the independent and interactive effects of the mean temperature and temporal autocorrelation (Fig. 5; AIC: 127.32, McFadden’s pseudo-*R*^2^ ≈ 0.46; see supplementary material S5). We found a significant negative effect of mean temperature on extinction risk across treatments (*p <* 0.001), as well as significant effects of brown noise and its interaction with mean temperature (*p <* 0.05), suggesting that high levels of autocorrelation modulate the relationship between mean temperature and extinction risk. Pink noise did not significantly change the extinction risk compared to white noise, and including neither initial population size nor tube number improved model fit. Both the weak and strong persistence boundaries were statistically significant predictors of extinction risk across all autocorrelation treatments (*χ*^2^ test, *p <* 0.0015), but they were strongest under pink noise conditions (*p <* 10^*−*5^). Tube location in the blocks (along the sides next to the heaters versus in the center farther from the heaters) was not a significant predictor of risk, supporting our assumption that heat diffused evenly across blocks.

Observed extinction risk across blocks followed the SSA model expectations relatively well (Fig. 6), but extinction risk was generally underpredicted (particularly under white noise conditions; Fig. 6a). We further compare the number of 12-hour time steps a population starting at *K* = 5000 individuals could survive at any given negative fitness value with the calculated 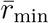 across Δ*t*s for each experimental time series (red line compared to other lines in Fig. 6d-f). This was an excellent predictor of risk for treatments whose 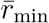 was near or below that boundary (W4-5; P3-5; B2-5), but risk was underpredicted for white noise treatments above the boundary.

**Figure 6:**
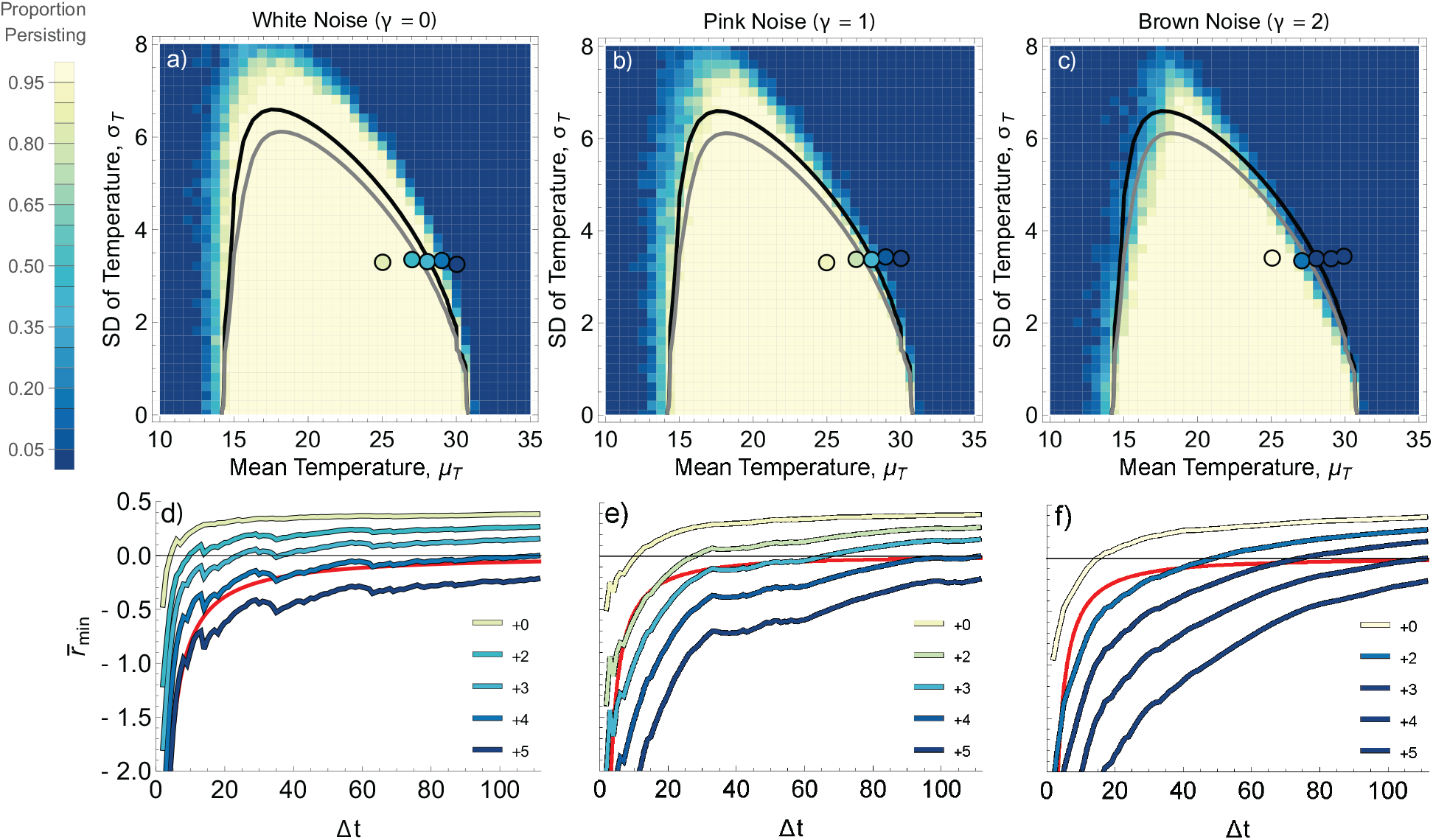
Experimental results (circles) compared to weak (gray) and strong (black) persistence boundaries, plotted over SSA model results averaged across 40 runs under the (a) white, (b) pink, and (c) brown noise temperature sequences. The minimum growth rates 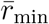 for time intervals of length Δ*t* for each mean temperature under the (d) white, (e) pink, and (f) brown noise temperature sequences, coloured by the proportion of extinct populations in the experiment, compared to the number of time steps the population is expected to persist with that negative growth rate from carrying capacity (red lines).

## 4 Discussion

Higher temporal autocorrelation increases extinction risk for populations with unimodal thermal tolerance. Over long time horizons, the autocorrelated *r*(*T*)-*α* model of logistic population growth shows a dramatic reduction in the envelope of thermal environments that permit population persistence. That envelope shrinks particularly quickly as noise shifts from white to pink (0.5 ≤*γ* ≤ 1), changes that natural atmospheric temperature time series have exhibited in recent decades (15), indicating that the ordering of temperatures through time may be an important facet of climate change to consider in risk forecasting. Results from our factorial experiment in *P. caudatum* microcosms confirmed that while increasing the mean temperature of variable time series increased extinction risk, how much it does so is modulated by the autocorrelation of that temperature time series. While the difference in extinction risk between low autocorrelation levels was not as stark in the experimental results as in the theoretical ones, we were able still able to predict the experienced extinction risk relatively well with a stochastic simulation algorithm and by comparing the minimum average fitness across the time series to the critical extinction boundary, calculated as the time to extinction from carrying capacity at a given negative fitness value. Across methods, increasing the autocorrelation of a time series increased extinction risk beyond what is predicted by the long-term average fitness or by the persistence boundaries defined by 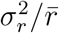 (although the latter remains a powerful predictor of risk at low levels of autocorrelation). Population dynamic models which can explicitly include temperature time series (and changes to their temporal structure) are an important tool for assessing whether the risks of unfavorable series of events are being overlooked, particularly in scenarios where the population is already experiencing a stressful thermal regime or temporal autocorrelation drastically increases.

Autocorrelation itself is not a universal driver of persistence versus extinction. While the broad theoretical debate of whether higher autocorrelation drives higher extinction risk was largely resolved by Schwager et al. (19), these conclusions have not been applied to population dynamic models where temperature has been incorporated in a biologically meaningful way. Our experimental results mainly demonstrate scenarios where higher autocorrelation is a disadvantage; the heated treatments under white and pink noise faced significantly less extinction risk than under brown noise, even though they experienced the exact same thermal distribution, simply because lower levels of autocorrelation allowed for unfavorable conditions to be spaced throughout the time series in a survivable sequence. However, there is some evidence that autocorrelation was advantageous under less stressful thermal regimes; the unheated brown noise populations experienced no extinctions (0, compared to 1 under pink noise and 2 under white), had the highest and least variable population densities (294±165, compared to 251±274 and 223±325 cells/mL), and potentially demonstrated some thermal acclimation in the post-experiment TPCs (see Fig. S7). When the temperature time series contained relatively few stressful temperatures, none of which were high above *T*_max_ (average of 1.7^*o*^, maximum of 3.2^*o*^), the smaller temperature changes that occur at each step in highly autocorrelated sequences may benefit the population by requiring organisms to acclimate less across time steps compared to pink and white noise.

Throughout this project, we have directly equated organismal body temperature with the temperature of the environment. Under thermally homogeneous experimental conditions, this is a very good assumption for free-living ectotherms like *P. caudatum*; however, real-world organisms experience heterogeneous environmental temperatures through the filters of thermoregulation and their own biophysical properties (e.g., 52; 53). How autocorrelation truly affects persistence is determined by the autocorrelation of the time series of an individual’s operative body temperature. In practice, this means that a more spatially homogeneous thermal environment (i.e., one with less opportunities for behavioral thermoregulation) may elevate extinction risk in the same way as a more autocorrelated temperature time series, because both conditions force organisms to spend longer periods of time under the stressful conditions that result in population decline (54).

In this paper, we have shown that predicting temperature-driven extinction risk without considering temporal autocorrelation overlooks an important interaction between population dynamics and the environment. Across stochastic simulations and a factorial experiment, higher autocorrelation corresponds to higher than expected extinction risk. That increase in risk is not driven by autocorrelation itself, but by the interaction between autocorrelation and thermal variation that exacerbates the negative effects of stressful temperatures by aggregating stressful events into sequences that drive extinction events. In thermal regimes that include stressful temperatures, autocorrelation thus increases the chances of experiencing deadly temperature sequences like heatwaves. Understanding this relationship is crucial for forecasting and managing for the long-term persistence of ectothermic organisms in a hotter, more variable climate.

