## Supplemental Material for "Temporal autocorrelation increases temperature-driven extinction risk by clustering stressful conditions"

**Title:** Temporal autocorrelation increases temperature-driven extinction risk
**Supplemental Material**

### S1 Additional Simulations

We simulated extinction risk across the parameter space using two different models, one using stochastic differential equations (SDE) and one using a stochastic simulation algorithm (SSA). The SDE model (described in Materials & Methods: Population Dynamic Modeling) is significantly faster and thus better suited to study the long-term behavior over the full parameter space; it can reasonably run for the length of time corresponding to the numerical approximation of the expected time to extinction at the weak boundary ($1/\alpha$ = 10,000 timesteps) over all mean, standard deviation, and autocorrelation treatments for a large number of replicates (100). Spectral synthesis is embedded within this model, such that each replicate generates its own time series with the desired level of autocorrelation.

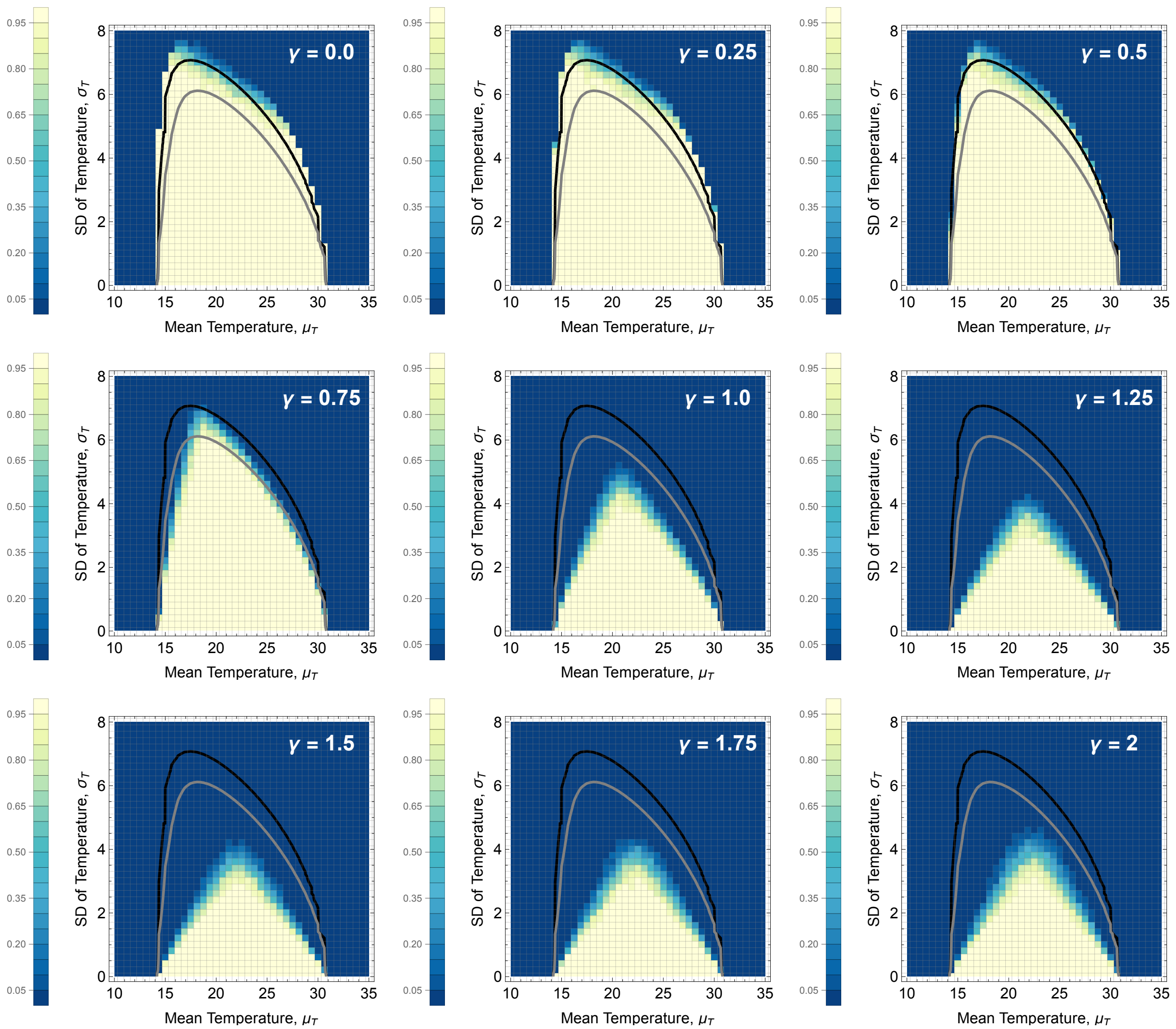

**Fig S1:** The simulated persistence envelope for additional values of the spectral exponent (γ) compared to the weak (gray) and strong (black) persistence boundaries using the SDE model. Diagonal panels (γ = 0.0, 1.0, 2.0) are also shown in Fig. 4a-c. Each parameter set was run 100 times for 10000 timesteps each.

The SSA model, on the other hand, is more computationally intensive and thus much slower, but easier to compare to the experimental results because it uses a different set of assumptions. The duration of each timestep $\delta_t$ can be set to the length used in the experimental treatments (12 hours, instead of an arbitrary 0.01 value) and, because a time series must be chosen prior to running the algorithm, we use the actual experimental temperature time series. The SSA model also incorporates demographic stochasticity. To transition from the SDE to the SSA model requires splitting the fitness TPC $r(T)$ into its component temperature-dependent birth and death rates (where $r(T) = b(T) - d(T)$). The density-dependent death rate increases linearly over time to reflect the deteriorating experimental conditions within microcosms. Over the course of the temperature time series, probabilistic birth and death events occur stochastically (with probabilities dependent on their temperature-dependent rates) at randomized time intervals according to Gillespie's algorithm (Gillespie 1977, Kummel & Vasseur 2025). Simulation results for 112 timesteps averaged across 40 replicates are shown in Fig. 6. Additional results for longer time periods (1009 timesteps, or 9 repeats of the experimental time series; averaged over 20 replicates) are shown in Fig. S2.

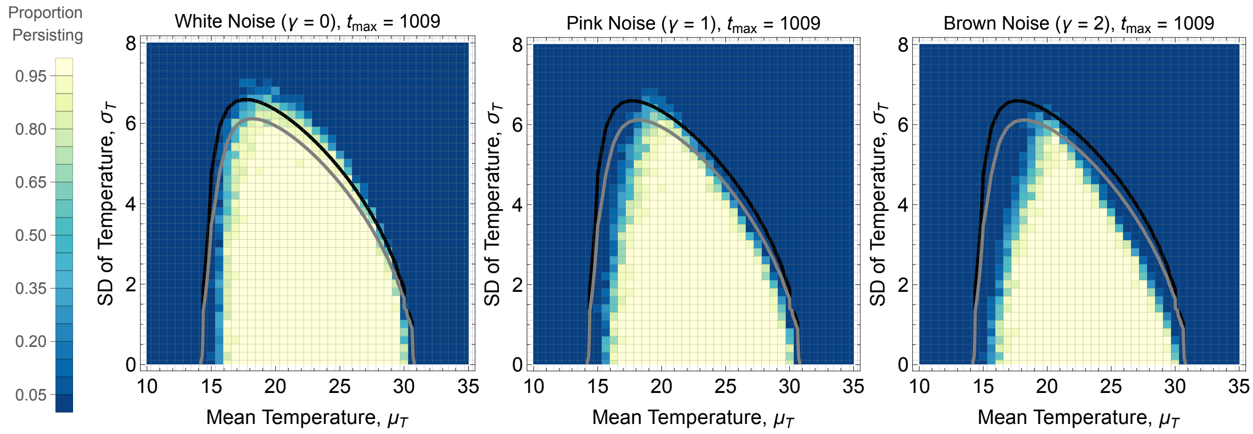


**Fig S2:** SSA results over 1009 timesteps (compared to 112 shown in Fig. 6), demonstrating the continued contraction of the persistence envelope as $t_{max}$ increases.

### S2 Temperature Probe Data

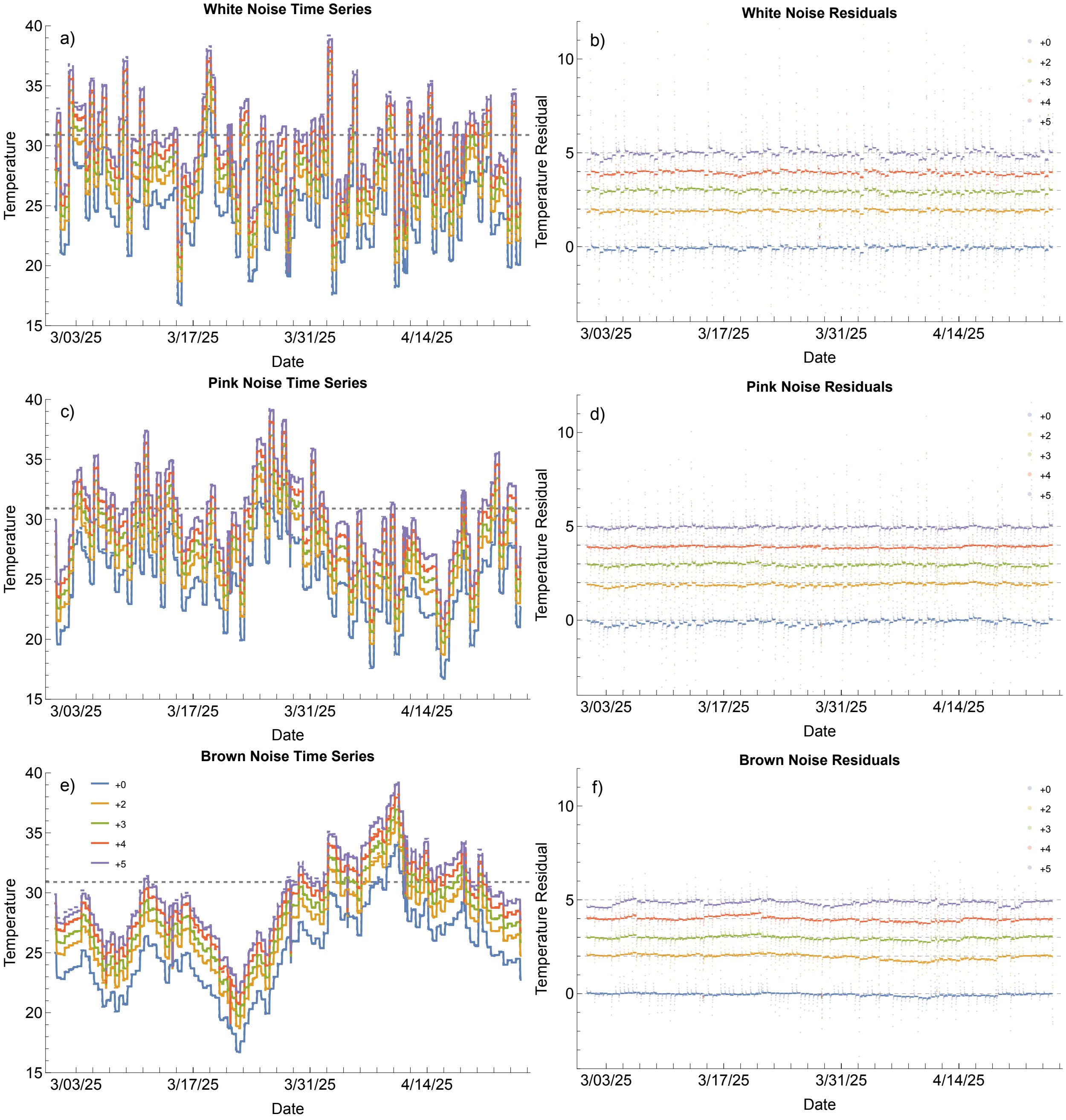


**Fig S3:** Comparison between goal and recorded temperature time series of each treatment, with left plots (a, c, e) showing the series themselves (dotted lines indicate goal temperatures, solid lines indicate recorded temperatures, straight dashed line indicates *P. caudatum*'s $T_{max}$) and right plots (b, d, f) showing temperature residuals (underlying dashed gray lines showing the intended offset for each block). The sporadic dots indicating high residuals in (d-f) show when the temperatures changed between time steps and when blocks were removed from incubators for medium replacement and counting. All recorded mean temperatures were within 0.05°C of goal temperatures, excluding the hottest brown noise treatment which ran approximately 0.1°C too cold.

| | Goal | | White | | Pink | | Brown | |
|---|---|---|---|---|---|---|---|---|
| | $\mu_T$ | $\sigma_T^2$ | $\mu_T$ | $\sigma_T^2$ | $\mu_T$ | $\sigma_T^2$ | $\mu_T$ | $\sigma_T^2$ |
| **+0** | 25 | 3.3 | 25.0093 | 3.2897 | 24.9601 | 3.2990 | 25.0382 | 3.3998 |
| **+2** | 27 | 3.3 | 26.9852 | 3.3502 | 26.9517 | 3.3783 | 27.0410 | 3.3421 |
| **+3** | 28 | 3.3 | 28.0213 | 3.3097 | 28.0088 | 3.3680 | 28.0284 | 3.3913 |
| **+4** | 29 | 3.3 | 28.9891 | 3.3356 | 28.9596 | 3.4265 | 29.0281 | 3.3898 |
| **+5** | 30 | 3.3 | 30.0071 | 3.2559 | 29.9975 | 3.3971 | 29.8907 | 3.4406 |

**Table S1:** Goal thermal distributions compared to observed thermal distributions (ºC).

| | Goal | | White | | Pink | | Brown | |
|---|---|---|---|---|---|---|---|---|
| | $\mu_T$ | $\sigma_T^2$ | $\mu_T$ | $\sigma_T^2$ | $\mu_T$ | $\sigma_T^2$ | $\mu_T$ | $\sigma_T^2$ |
| **+0** | 25 | 3.3 | 0.0093 | -0.0103 | -0.0399 | -0.0009 | 0.0382 | 0.0998 |
| **+2** | 27 | 3.3 | -0.0148 | 0.0502 | -0.0483 | 0.0783 | 0.0410 | 0.0421 |
| **+3** | 28 | 3.3 | 0.0213 | 0.0097 | 0.0088 | 0.0680 | 0.0284 | 0.0913 |
| **+4** | 29 | 3.3 | -0.0109 | 0.0356 | -0.0404 | 0.1265 | 0.0281 | 0.0898 |
| **+5** | 30 | 3.3 | 0.0071 | -0.0441 | -0.0025 | 0.0971 | -0.1093 | 0.1406 |

**Table S2:** Residuals of goal thermal distributions compared to observed thermal distributions (ºC).

### S3 Full Experimental Results

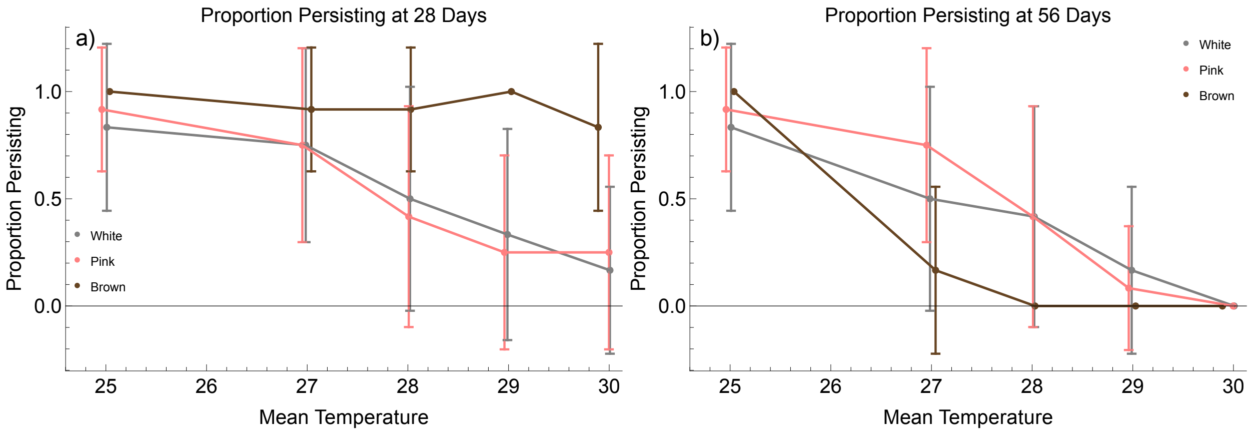


**Fig S4:** Proportion of replicates persisting at the halfway (a) and final (b) cell counts (error bars show standard deviation).

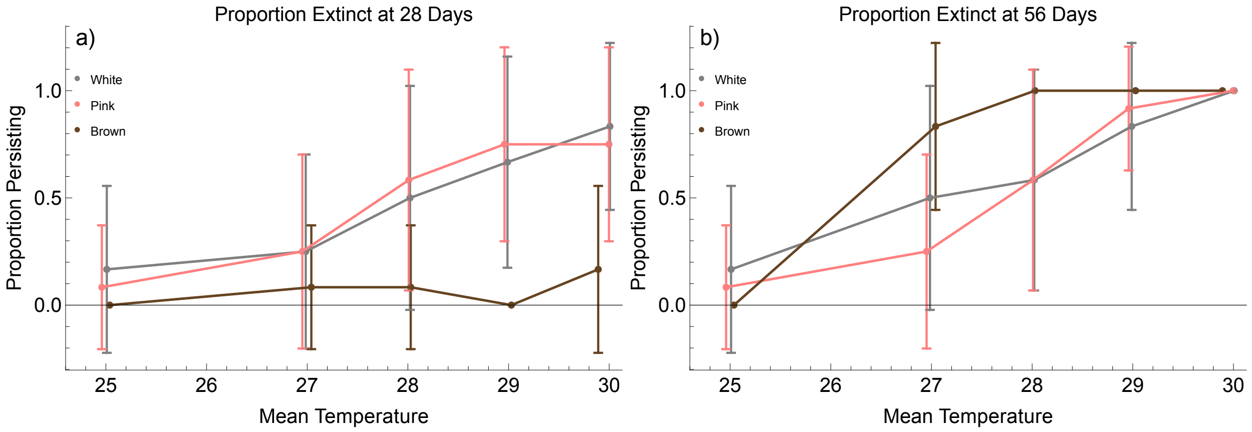


**Fig S5:** Proportion of replicates extinct at the halfway (a) and final (b) cell counts (error bars show standard deviation).

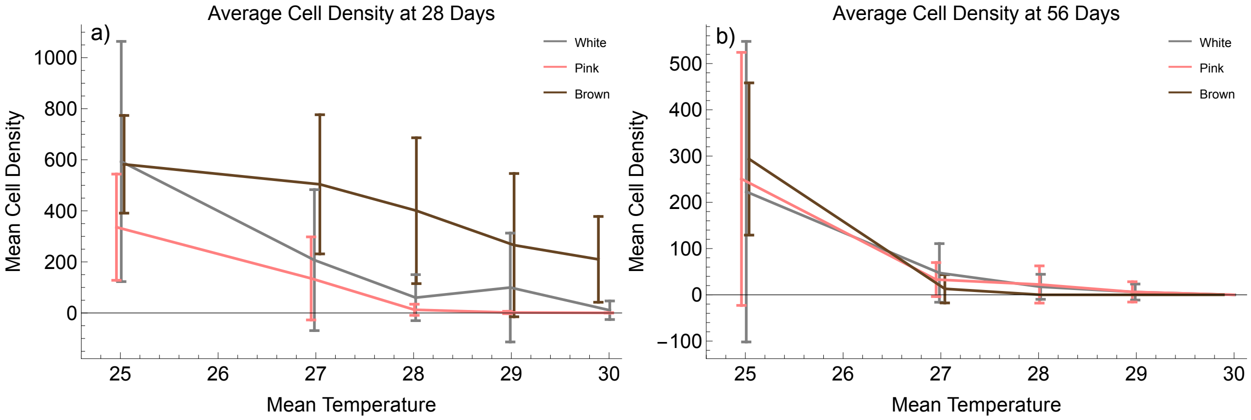


**Fig S6:** Average cell densities at the halfway (a) and final (b) time cell counts (error bars show standard deviation).

### S4 TPC Data

We gathered thermal performance curve (TPC) data for the protist *P. caudatum* using the methodology of Wieczynski et al. (2021) and described in Materials and Methods: Thermal Performance Curves. The pre-experiment TPC is the same one gathered for and used in Vasseur et al. (2025), collected in the summer of 2024. The post-experiment TPCs were collected two days prior to the end of the experimental run. The reduced thermal tolerance of the post-experiment stock culture compared to the pre-experiment stock culture is likely due to the shorter time period over which TPCs were collected, as well as the reduced number of replicates and different temperatures used; TPCs are best compared across equivalent conditions (e.g., within those shown in Fig. S6, but not between Fig. S6 and Fig. 2a). We fit the 'lactin2' model (Lactin et al., 1995) with the 'rTPC' package (Padfield et al., 2021) in *R* v.4.4.3; the lactin2 parameters are $a$ (the steepness of the rising portion of a TPC), $b$ (the TPC height), $T_{max1}$ (temperature at which curve decelerates beyond $T_{opt}$; usually written as $T_{max}$), and $\delta_T$ (the thermal safety margin).

| Parameter | Estimate | SE | T-statistic | 95% CI low | 95% CI high | p-value |
|---|---|---|---|---|---|---|
| $a$ | 0.044 | 0.0373 | 1.18 | -0.0304 | 0.118 | 0.24 |
| $b$ | -1.77 | 0.613 | -2.89 | -2.998 | -0.551 | 0.0051 |
| $T_{max1}$ | 35.3 | 3.34 | 10.56 | 28.594 | 41.913 | 5.52e-16 |
| $\delta_T$ | 5.43 | 6.41 | 0.85 | -7.353 | 18.222 | 0.40 |

**Table S3** Pre-experiment TPCs: Parameter estimates for 'lactin2' TPC model fit in *P. caudatum*, based on 12 data points per temperature at 6 temperatures (18, 22, 24, 26, 28, 30). Plot of TPC and experimental data can be seen in Fig. 2c.

| *Source* | Parameter | Estimate | SE | T-statistic | 95% CI low | 95% CI high | p-value |
|---|---|---|---|---|---|---|---|
| Stock | $a$ | 0.0878 | 7.17 | 0.012 | -14.91 | 15.0 | 0.990 |
| | $b$ | -3.25 | 97.7 | -0.033 | -207.69 | 201.20 | 0.974 |
| | $T_{max1}$ | 33.3 | 84.1 | 0.396 | -142.68 | 209.32 | 0.696 |
| | $\delta_T$ | 6.89 | 356 | 0.019 | -738.74 | 752.54 | 0.985 |
| W1-10 | $a$ | 0.0799 | 7.19 | 0.011 | -14.92 | 15.08 | 0.991 |
| | $b$ | -2.99 | 97.4 | -0.031 | -206.15 | 200.16 | 0.976 |
| | $T_{max1}$ | 35.2 | 89.1 | 0.395 | -150.72 | 221.12 | 0.697 |
| | $\delta_T$ | 7.39 | 435 | 0.017 | -900.53 | 915.33 | 0.987 |
| W4-6 | $a$ | 0.0215 | 0.012 | 1.817 | -14.92 | 15.08 | 0.088 |
| | $b$ | -1.76 | 0.409 | -4.298 | -206.15 | 200.16 | 5.54e-4 |
| | $T_{max1}$ | 31.5 | 1.61 | 19.55 | -150.72 | 221.12 | 1.36e-12 |
| | $\delta_T$ | 1.19 | 1.33 | 0.894 | -900.53 | 915.33 | 0.384 |
| P1-9 | $a$ | 0.0197 | 0.014 | 1.397 | -0.0102 | 0.049 | 0.1814 |
| | $b$ | -1.61 | 0.498 | -3.242 | -2.668 | -0.558 | 0.0051 |
| | $T_{max1}$ | 30.69 | 1.726 | 17.78 | 27.029 | 34.35 | 5.83e-12 |
| | $\delta_T$ | 0.664 | 1.650 | 0.402 | -2.834 | 4.162 | 0.6927 |
| P4-2 | $a$ | 0.0858 | 1.98 | 0.043 | -4.045 | 4.217 | 0.966 |
| | $b$ | -3.45 | 25.54 | -0.135 | -56.72 | 49.82 | 0.894 |
| | $T_{max1}$ | 34.02 | 19.92 | 1.707 | -7.55 | 75.58 | 0.103 |
| | $\delta_T$ | 6.95 | 107.8 | 0.064 | -217.98 | 231.9 | 0.949 |
| B1-6 | $a$ | 0.0790 | 6.76 | 0.012 | -14.03 | 14.19 | 0.991 |
| | $b$ | -2.544 | 92.88 | -0.027 | -196.3 | 191.2 | 0.978 |

| | | | | | | | |
|---|---|---|---|---|---|---|---|
| | $T_{max1}$ | 35.23 | 90.83 | 0.388 | -154.2 | 224.7 | 0.702 |
| | $\delta_T$ | 7.482 | 410.0 | 0.018 | -947.8 | 862.8 | 0.986 |

**Table S4** Post-experiment TPCs: Parameter estimates for 'lactin2' TPC model fit in *P. caudatum*, based on 4 data points per temperature at 6 temperatures (18, 20, 22, 26, 28, 30). The chosen replicates were the stock culture (a), which had been in the constant temperature incubator at a 25°C; the highest density persisting replicates in each of the lowest mean temperature treatments W1 (b), P1 (d), and B1 (f), to compare any baseline changes across autocorrelation levels; and the highest density replicates in the hottest treatment to persist across both white and pink noise W4 (c) and P4 (e), to see whether the replicate persisting in P4 had survived due to significant changes to thermal tolerance. The limited number of data points resulted in low-confidence parameter estimates across the board, so resulting TPCs should be interpreted with caution.

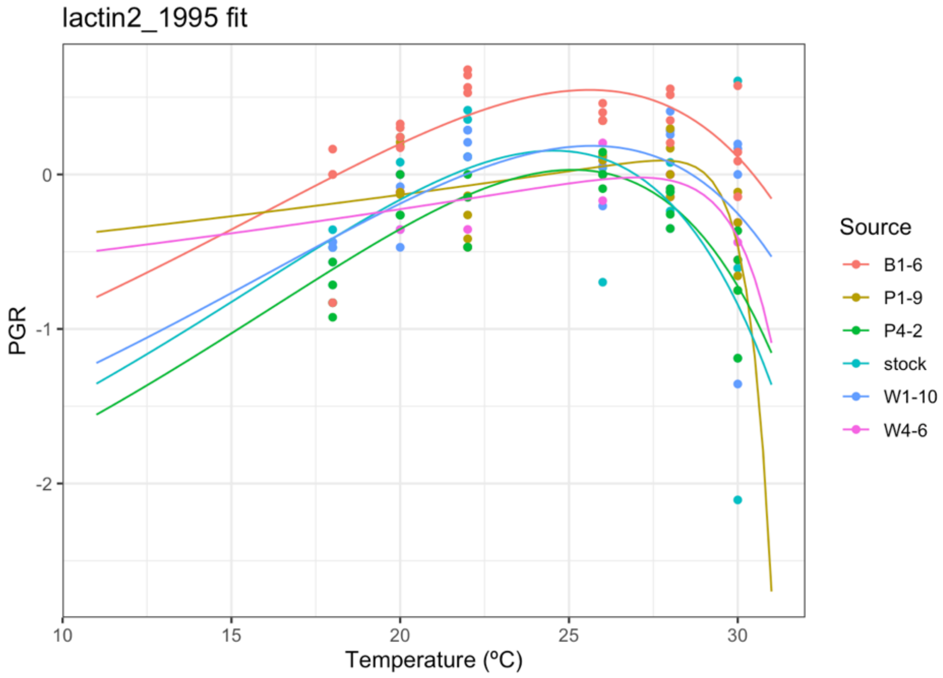


**Fig S7:** 'lactin2' TPC model fits of population growth rates (PGR) per day for post-experimental *P. caudatum* TPCs (lines) plotted with experimental points (dots).

### S5 Statistical Model Fitting

We fit our data using bias-reduced binary generalized linear models. We used bias-reduction GLMs ('brglm' package; Kosmidis, 2021) instead of standard GLMs ('glm' package) because of data separation. We treated mean temperature as a continuous variable and autocorrelation of temperature as a categorical variable to reflect the experimental setup. The best fit model included the main effects and interaction effects of the mean and autocorrelation of temperature (Extinction ~ Mu * Gamma; Table S5). We rejected models including the initial population density (as main effect: AIC 129.42, same terms significant as chosen model; as interactive effect: AIC 132.62, no significant terms found) and the tube number (as main effect: AIC 129.35, same terms significant as chosen model; as interactive effect: AIC 133.45, no significant terms found) because the added terms were never statistically significant and always increased AIC values and model complexity without improving the model fit. We additional tested binary predictors with $X^2$ tests (Tables S6, S7, S8). All tests were done in *R* v4.4.3.

| | **Estimate** | **Std Error** | **z-value** | **Pr(>\|z\|)** | **Signif** | **2.5% CI** | **97.5% CI** |
|---|---|---|---|---|---|---|---|
| **Intercept** | 22.6673 | 6.1452 | 3.689 | 0.000225 | *** | 10.62 | 34.71 |
| **Mu** | -0.8374 | 0.2226 | -3.761 | 0.000169 | *** | -1.274 | -0.4010 |
| **Gamma1** | 11.7243 | 10.7144 | 1.094 | 0.273842 | | -9.275 | 32.72 |
| **Gamma2** | 42.6357 | 21.6028 | 1.974 | 0.048425 | * | 0.2949 | 84.98 |
| **Mu:Gamma1** | -0.4122 | 0.3863 | -1.067 | 0.285969 | | -1.169 | 0.3449 |
| **Mu:Gamma2** | -1.6339 | 0.8026 | -2.036 | 0.041790 | * | -3.207 | -0.0607 |

**Table S5:** Summary of the bias-reduction generalized linear model testing for the effects of experimental parameters on extinction risk in *P. caudatum.* The best fit model included measured mean temperature (represented by the continuous variable Mu), autocorrelation of temperature (represented by the categorical variable Gamma, where Gamma1 indicates pink noise and Gamma2 indicates brown noise), and a binary response variable (0 = extinct, 1 = persisting). This model had an AIC of 127.32, explained 45.66% of deviance compared to the null (McFadden's pseudo-$R^2$ = 1 – model deviance/null deviance), and there was no indication of overdispersion (ratio of residual deviance to residual degrees of freedom = 0.66). For this model, null, residual, and penalized deviance were 212.24 (on 179 degrees of freedom), 115.32 (on 174 degrees of freedom), and 104.3053, respectively. 0.001 significance level is indicated by *** and 0.05 level by *.

| | **Edge** (1-4, 9-12) | **Middle** (5-8) | |
|---|---|---|---|
| **Persisting** | 40 | 23 | *63* |
| **Extinct** | 80 | 37 | *117* |
| | *120* | *60* | |

**Table S6** Contingency table comparing extinction risk on the edge versus middle of block ($X^2$ = 0.24725, p-value = 0.619).

| | **Inside** | **Outside** | |
|---|---|---|---|
| a) *All Treatments – Strong Boundary* | | | |
| **Persisting** | 60 | 3 | *63* |
| **Extinct** | 48 | 69 | *117* |
| | *108* | *72* | |

| b) *All Treatments – Weak Boundary* | | | |
|---|---|---|---|
| **Persisting** | 50 | 13 | *63* |
| **Extinct** | 22 | 95 | *117* |
| | *72* | *108* | |
| c) *White Noise – Strong Boundary* | | | |
| **Persisting** | 21 | 2 | *23* |
| **Extinct** | 15 | 22 | *37* |
| | *36* | *24* | |
| d) *White Noise – Weak Boundary* | | | |
| **Persisting** | 16 | 7 | *23* |
| **Extinct** | 8 | 29 | *37* |
| | *24* | *36* | |
| e) *Pink Noise – Strong Boundary* | | | |
| **Persisting** | 25 | 1 | *26* |
| **Extinct** | 11 | 23 | *34* |
| | *36* | *24* | |
| f) *Pink Noise – Weak Boundary* | | | |
| **Persisting** | 20 | 6 | *26* |
| **Extinct** | 4 | 30 | *34* |
| | *24* | *36* | |
| g) *Brown Noise – Strong Boundary* | | | |
| **Persisting** | 14 | 0 | *14* |
| **Extinct** | 22 | 24 | *46* |
| | *36* | *24* | |
| h) *Brown Noise – Weak Boundary* | | | |
| **Persisting** | 14 | 0 | *14* |
| **Extinct** | 10 | 36 | *46* |
| | *24* | *36* | |

**Table S7** Contingency tables comparing extinction risk inside versus outside the strong (a, c, e, g) and weak (b, d, f, h) extinction boundaries across all autocorrelation treatments (a, b) and within each autocorrelation treatment (White: c-d, Pink: e-f, Brown: g-h)

| | **Weak Boundary** | | **Strong Boundary** | |
|---|---|---|---|---|
| | $X^2$ | p-value | $X^2$ | p-value |
| **All** | 60.082 | $9.097\times10^{-15}$ | 47.913 | $4.455\times10^{-12}$ |
| **White** | 11.66 | $2.818\times10^{-4}$ | 11.66 | $6.386\times10^{-4}$ |
| **Pink** | 23.419 | $1.303\times10^{-6}$ | 22.401 | $2.213\times10^{-6}$ |
| **Brown** | 24.227 | $8.56\times10^{-7}$ | 10.097 | $1.485\times10^{-3}$ |

**Table S8** Summary of significance of $X^2$ tests. All p-values indicate that extinction is statistically more significant outside both extinction boundaries across all treatments.

### S6 Pilot Experiment Results

We conducted a pilot version of the experiments presented in this paper from same *P. caudatum* culture. This experiment followed the same methodology, but included 8 mean temperatures (20, 23, 26, 27, 28, 29, 30, 31) instead of 5 (25, 27, 28, 29, 30) and 2 autocorrelation treatments (white and pink) instead of 3 (white, pink, and brown); different temperature time series were used. Results, though not inconsistent with the results of the final experiment, were discarded due to much poorer monitoring and control of the block temperatures, which resulted in poor comparisons of mean temperature treatments between autocorrelation treatments. However, they were used to guide the choice of mean temperature treatments for the final experiments.

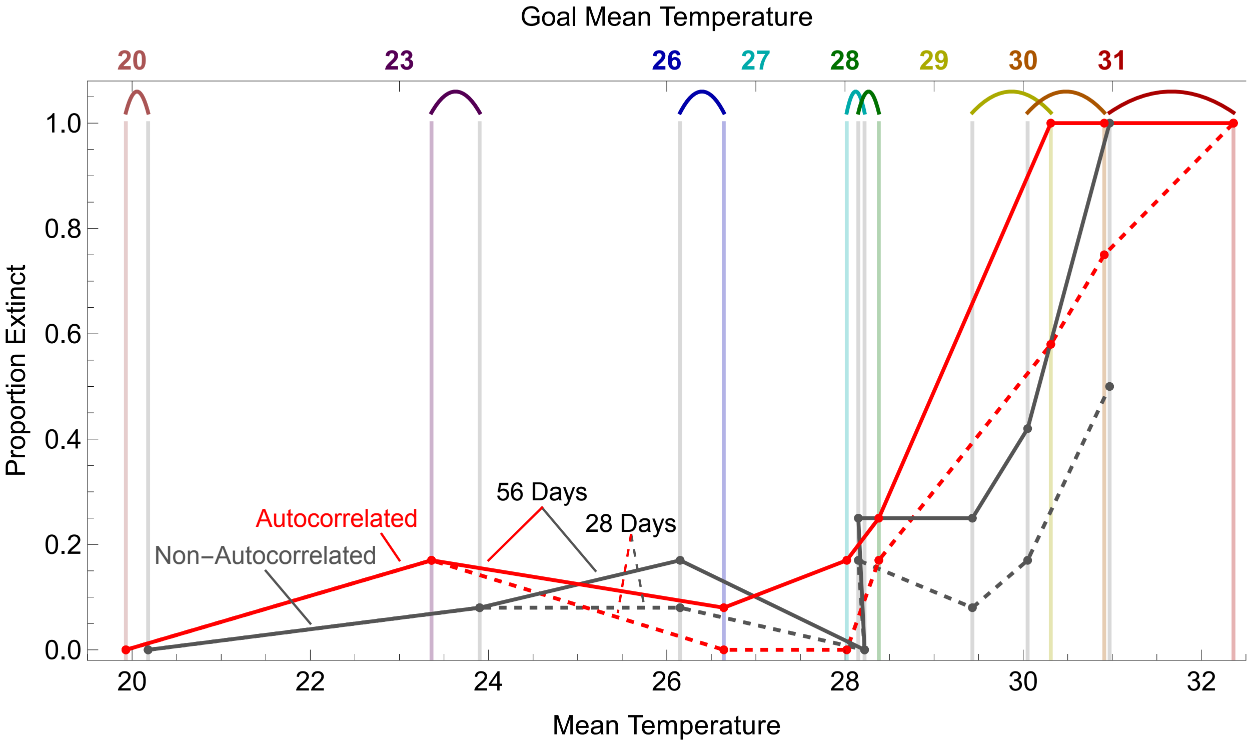


**Fig S8:** Pilot results after 28 (dotted lines) and 56 (solid lines) days comparing the proportion of populations to go extinct at each mean temperature experiencing either an autocorrelated (red) or non-autocorrelated (gray) time series. Individual treatments, represented by points, are plotted at the actual mean temperature experienced by that block; the goal temperatures are shown at the top. Gaps between the experienced temperatures of blocks intended to be at the same mean temperature are shown with colored brackets.
